# Intratumoral microbiota-derived S1P sensitizes the combination therapy of Capecitabine and Nivolumab

**DOI:** 10.1101/2025.06.05.658030

**Authors:** Chen-Shu Dai, Sen-Tao Pan, Tian-Tian Qi, Hui-Ling Shang, Ri-Hua Xie, Hao Liu, Zhen-Ming Liu, Yi-Min Cui, Yu-Hang Zhang

**Author notes:** These authors contributed equally to this work (Chen-Shu Dai, Sen-Tao Pan, Tian-Tian Qi). Corresponding authors: Professors Yu-Hang Zhang, Yi-Min Cui and Zhen-Ming Liu. **Author’s contribution:** Yu-Hang Zhang, Yi-Min Cui and Zhen-Ming Liu have conceived the project. Hui-Ling Shang and Ri-Hua Xie collected clinical samples and pathological analysis. Sen-Tao Pan and Chen-Shu Dai have implemented experiments of pharmaceutical analysis. Tian-Tian Qi and Hao Liu performed statistical analysis for multi-omics data. Chen-Shu Dai and Sen-Tao Pan wrote the manuscript, which was edited by all authors.

## Abstract

Clinical responses of colorectal cancer (CRC) treatments vary considerably due to the heterogeneity of tumor micro-environment (TME), where intratumoral microbiota may reshape the unique inflammation imprints. However, its complex mechanistic underpinnings remain incompletely elucidated. Herein we sought to delineate the critical role of intratumoral microbiota in potentiating combination therapeutics against CRC. By comparing germ-free (GF) and specific pathogen-free (SPF) mouse models of 33 potential CRC treatments, we screened out the combination regimen of Capecitabine-Nivolumab significantly augmented by intratumoral microbiota in tumor regression and survival prolongation. The enrichment of *Bacteroides fragilis* induced by Capecitabine-Nivolumab was concomitant with elevated microbial sphingosine-1-phosphate (S1P), which further upregulated tumoral PD-L1 expression by enhancing histone deacetylation at the *CD274* locus. Activation of microbial sphingosine kinase 2 (SphK2) ultimately led to an expansion of effector memory *CD8^+^* T cells and reduction of exhausted T cell subsets within TME. To conclude, these findings advance our understanding of the intricate interplay among intratumoral microbiota, sphingolipid metabolites and immunochemotherapeutics. It highlights microbial sphingolipids as potential predictive biomarkers for strategies of targeting intratumoral microbiota in CRC management.

## Introduction

Colorectal cancer (CRC) has been reckoned as one of the most common malignant tumor types worldwide, with the incidence and mortality rates increasing annually.(Bray *et al*, 2024; Morgan *et al*, 2023) As the research into its molecular mechanisms and therapeutic strategies deepened, an ever-growing number of combination regimens, including chemotherapy, targeted therapies and immune checkpoint inhibitors, have been introduced into clinical practice for CRC patients.(Li *et al*, 2024; Li *et al*, 2025; Van Cutsem *et al*, 2014) But owing to individual factors and tumor heterogeneity, the same regimen often demonstrated markedly different clinical benefits across CRC patients, underscoring the urgent need to identify more effective and broadly adaptable combination therapies.(Boshuizen & Peeper, 2020; Shin *et al*, 2023) In recent years, the role of intratumoral microbiome has garnered increasing attention in the onset, progression and treatment of CRC. Mounting evidence suggested that the composition and diversity of intratumoral microbiota not only influenced the tumor immune micro-environment and disease progression, but also closely correlated with drug efficacy and adverse reactions.(Bullman, 2023; Lu *et al*, 2024; Triner *et al*, 2019; Yang *et al*, 2023; Yang *et al*, 2017) Accordingly, how to effectively incorporate intratumoral microbiota into the assessment of therapeutic efficacy necessitates systematic investigation.

To better elucidate how intratumoral microbiome mediates these treatments, we developed a screening platform for various CRC therapies. By comparing the tumor number and tumor volume between germ-free (GF) *versus* specific pathogen-free (SPF) mice, we evaluated the efficacy differences for all potential combination approaches. Among these therapies we tested, four regimens indicated the most significant discrepancies in therapeutic efficacy, including the combined use of Capecitabine and Nivolumab. Capecitabine is a fluoropyrimidine pro-drug and remains the only globally approved agent that can be administered orally at home.(Bando *et al*, 2023) It was synthesized in the 1990s,(Malet-Martino & Martino, 2002) which received approval from the U.S. Food and Drug Administration (FDA) in 2005.(Vodenkova *et al*, 2020) As the monotherapy regimen of fluoropyrimidines, Capecitabine is the first-choice treatment for metastatic CRC in the first-line setting.(Mettu *et al*, 2022; Twelves *et al*, 2001) Nivolumab, a highly selective human IgG4 monoclonal antibody to inhibit the programmed death receptor-1 (PD-1), is frequently applied in patients with mismatch repair-deficient/microsatellite instability-high (dMMR/MSI-H) metastatic CRC (mCRC).(Parikh *et al*, 2021; Sidaway, 2024) Although both agents displayed considerable anti-tumor effects, their combination therapy has not yet been adopted in clinical practice, whose underlying mechanisms and therapeutic potentials remain to be investigated.

In this study, the overarching aim was to elucidate the complex host-microbiome interactions and intratumoral mechanisms underpinning the combined treatment of Capecitabine and Nivolumab, paving new avenues to improve this regimen in the colorectal environment. Identifying specific intratumoral microbial strains that undergo notable changes under the combination therapy, we conducted metabolomic analyses to differ microbial metabolites across treatment groups. Subsequently, we integrated epigenetic profiling to determine whether these microbial metabolites affected the downstream gene expression of CRC progression. In light of Nivolumab being an immune checkpoint inhibitor, we performed single-cell sequencing to compare immunological variations among groups, comprehensively exploring the potential way to enhance therapeutic efficacy of Capecitabine and Nivolumab combination therapy.

## Results

### Screening of CRC treatment regimens significantly affected by intratumoral microbiota

To establish a screening platform for 33 different treatment regimens of CRC patients, we conducted experiments on C57BL/6 mice raised by germ-free (GF) and conventional (specific pathogen-free, SPF) conditions. Using azoxymethane (AOM)/dextran sulfate sodium (DSS) to construct CRC model, we administered different cancer therapies and collected CRC tissues from each group for metagenome and metabolome analyses (Fig. 1A). Among 33 microbial communities validated *in vivo*, we identified four regimens with the highest Z-scores: Capecitabine combined with Nivolumab, Capecitabine combined with Irinotecan and Bevacizumab, CAPEFOX combined with Irinotecan, CAPEFOX alone (Fig. 1B). Fig. 1C manifests the effects of these four treatment regimens on tumor growth between SPF and GF mice. It revealed that drug efficacy was more pronounced in SPF mice, with the greatest difference observed under Capecitabine-Nivolumab combination therapy. Upon this screening, we initially identified the interactions between 33 treatment regimens and 41 representative microbial strains, from which we selected 39 potential bacterium-therapy interaction pairs. These pairs underwent two rounds of independent validations (Fig. 1D). We set a false discovery rate (FDR)-adjusted *P* value < 0.05 and a drug consumption rate of over 30% as the thresholds. These targeted interactions were further validated in GF mice with colonized bacteria, revealing an *in vivo* interaction network that covered all tested strains and approximately 36.36% of therapies (12 out of 33 therapies).

**Figure 1.**
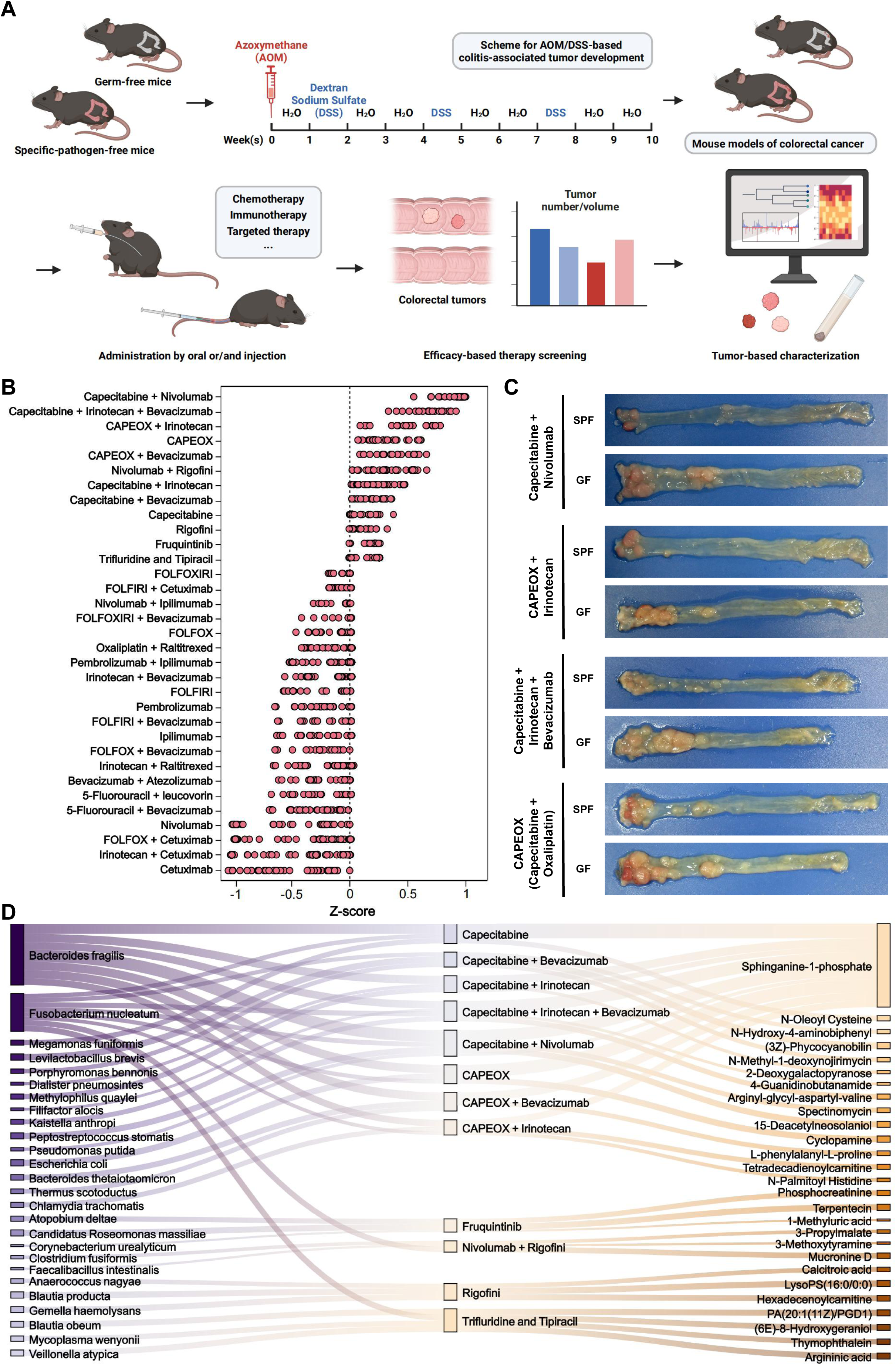
Comparative screening of anti-tumor efficacy between SPF and GF mice revealed the role of intratumoral microbiota in Capecitabine-Nivolumab combination. **(A)** Schematic workflow of intratumoral microbiota-mediated anti-tumor efficacy screening platform, which was created by Biorender. Age-matched SPF and GF mice were simultaneously treated with AOM/DSS for 10 weeks, prior to 4 weeks of anti-tumor treatments. For each group of mice, theraputic efficacy was evaluated by tumor number and volume, and CRC tissues were collected for microbiological detection. **(B)** *Z* factor of intratumoral microbiota effect on the given therapies in each SPF-GF mice pair, *n* = 6_SPF_ * 6_GF_ = 36. The value of *Z* factor > 0.5 indicates the positive effect. **(C)** Colorectal longitudinal sections of SPF and GF mice treated with combinations of Capecitabine-Nivolumab, CAPEOX (Capecitabine and Oxaliplatin), CAPEOX-Irinotecan, CAPEOX-Irinotecan-Bevacizumab for 4 weeks. **(D)** Microbiota-therapy-metabolite interaction network identified in this study. Left network: Effects of intratumoral bacteria on theraputic efficacy. Significant interactions in two independent screenings (*n* = 3 per screen) were validated in a follow-up assay (*n* = 3; FDR-corrected *P* < 0.05) are shown (Spearman’s rank tests). Right network: differential intratumoral metabolites between SPF and GF mice of theraputic efficacy detected by two independent screenings (Spearman’s rank tests).

### The enrichment of intratumoral *Bacteroides fragilis* promoted the combination treatment of Capecitabine and Nivolumab in CRC

To further investigate the mechanisms underlying the differential efficacy of Capecitabine-Nivolumab combination therapy between SPF and GF mice, we randomly assigned all SPF and GF models of colorectal cancer into four treatment groups: saline control, Capecitabine alone, Nivolumab alone, combination of Capecitabine with Nivolumab. During this period, we recorded the body weight changes, diarrhea scores and fecal blood scores for each group. Compared with GF mice, SPF mice receiving the Capecitabine-Nivolumab combination maintained more stable body weights and exhibited significantly lower diarrhea and fecal blood scores (Figs. 2A and EV1A,B). After sacrifice, SPF mice of Capecitabine-Nivolumab combination group showcased a pronounced reduction in colorectal tumor size. By contrast, GF mice demonstrated some degrees of tumor shrinkage but remained with larger tumor volumes, suggesting a potential role of microbial environment in treatment efficacy (*P* < 0.01; Figs. 2B-D and EV1C). To further assess how different CRC treatment regimens reversely affected intratumoral microbiota, we collected CRC tissues from SPF mice for metagenomic sequencing, obtaining an average of 45,832,565 sequencing reads per sample. Removing unannotated species shown in MetOrigin database (https://data.mendeley.com/drafts/55h4fxt79t), we identified 394 microbial species classified into 157 families and 271 genera. According to α-diversity indices (ACE, Chao1, Shannon, Simpson), no significant discrepancies were found in microbial richness or diversity among the groups (Fig. EV1D). PCA analysis (*R*² = 0.17, *P* = 0.016) and PCoA analysis (*R*² = 0.71, *P* = 0.027) indicated their variations in β-diversity features (Figs. 2E and EV1E). NMDS analysis demonstrated a clustering trend across groups (Stress = 0.175, Fig. EV1E). Subsequently, we examined the taxonomic composition of intratumoral microbiota in SPF mice (Fig. 2F). Through ultra-deep metagenomic sequencing and LEfSe analysis, we determined that *Bacteroides fragilis* (*B. fragilis*) was prominently enriched in both Capecitabine group and Capecitabine-Nivolumab combination group (Fig. 2G,H). This enrichment was further confirmed by RT-qPCR in SPF mice treated with Capecitabine alone or in combination with Nivolumab (Fig. 2I). Intratumoral microbial co-occurrence network illustrated the dominating microbes associated with *B. fragilis* (Fig. EV1F), suggesting that *B. fragilis* inevitably maintained the stability of intratumoral microbiome composition during Capecitabine-Nivolumab combination therapy.

**Figure 2.**
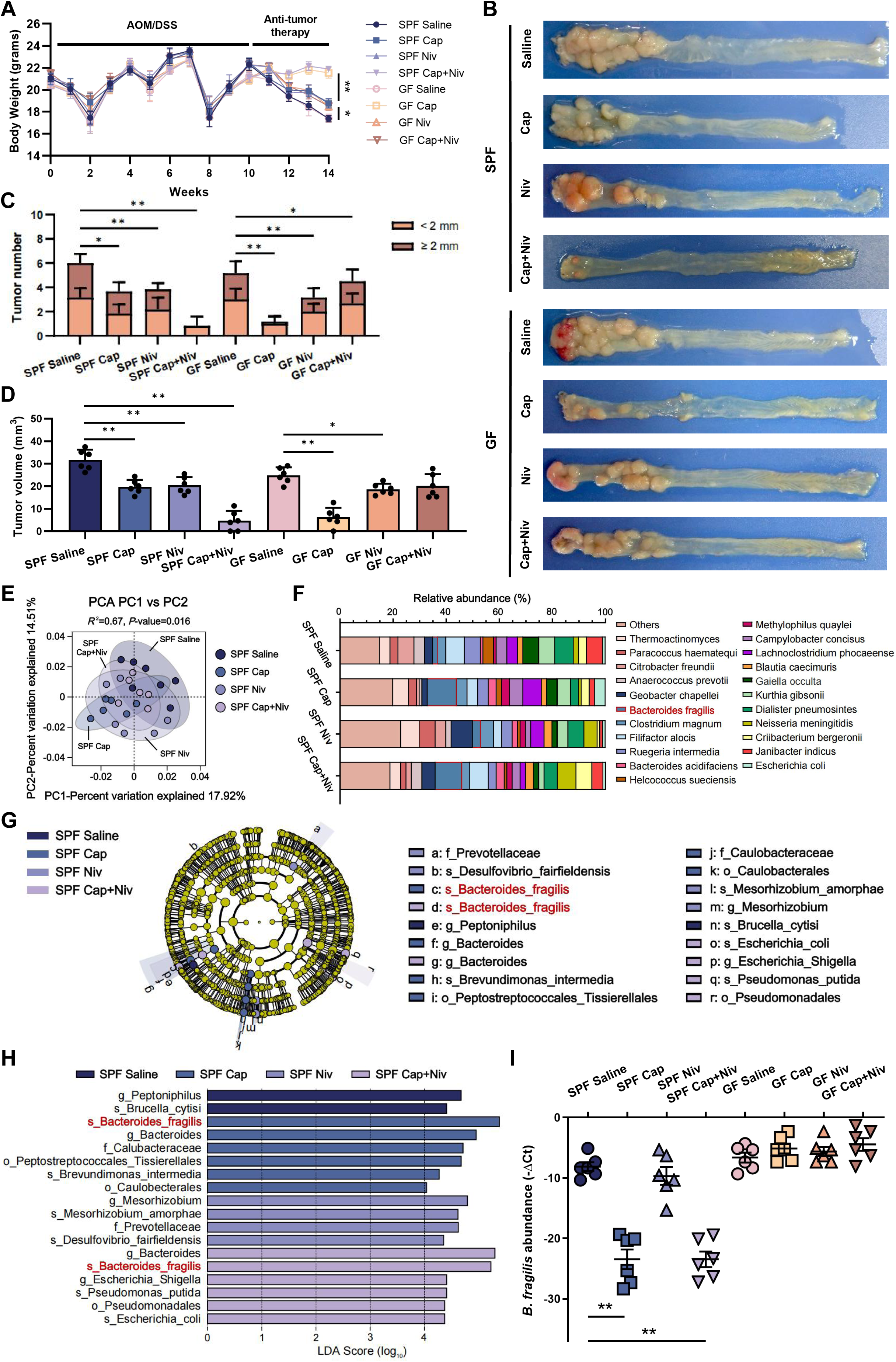
*B. fragilis* was enriched by the treatment of Capecitabine. **(A)** Body weights of SPF and GF mice in each group during AOM/DSS induction and treatments (saline, Capecitabine, Nivolumab, Capecitabine-Nivolumab combination). *n* = 6 mice per group (the same applies hereinafter). Data are means ± *SD*, \**P* < 0.05, \*\**P* < 0.01 (Student’s *t*-tests). **(B)** Colorectal longitudinal sections of each group after 4-week anti-tumor therapies. **(C,D)** Colorectal tumor number **(C)** and volume **(D)** across groups. Data are means ± *SD*, \**P* < 0.05, \*\**P* < 0.01 (Student’s *t*-tests). **(E)** Principal coordinate analysis (PCA) scatter-plot of bacterial community β-diversity based on Bray-Curtis distances among the SPF mice groups after treatments. **(F)** Species taxa summary of the altered intratumoral microbiome in SPF mice treated with Capecitabine or/and Nivolumab. **(G,H)** Linear discriminant analysis (LDA) measurements to identify the significant abundance of intratumoral microbiota. Taxonomic cladogram **(G)** and histogram **(H)** were generated by LDA Effect Size (LEfSe) among groups. Only LDA scores > 4 are shown. **(I)** RT-qPCR to determine the 16S rRNA segment of *B. fragilis* absolute abundance in each group. Data are means ± *SD*, \**P* < 0.05, \*\**P* < 0.01 (Student’s *t*-tests).

### *B. fragilis*-derived sphingosine-1-phosphate upgraded intratumoral PD-L1 levels by inhibiting HDAC1 activity

Given the potential mechanism underlying the efficacy of intratumoral microbiota metabolism on synergistic effect of Capecitabine and Nivolumab, we sought to elucidate the metabolic alterations of inter-cellular microecology between GF and SPF mice. Metabolome datasets of the volcanic plots showcased higher sphingosine-1-phosphate (S1P) among all substantially discrepant intratumoral metabolites triggered by the combination treatment of Capecitabine-Nivolumab in SPF mice, comparing with SPF mice treated with Saline as well as GF mice treated with Capecitabine-Nivolumab combination (Fig. 3A). Neither Capecitabine alone nor Nivolumab alone had obviously affected intratumoral microbiota-derived S1P for both SPF and GF mice (Fig. EV2A). KEGG enrichment of these multiple differences were mainly concentrated on sphingolipids (SLs) metabolism (9.52%, Fig. EV2B). By employing ultra performance liquid chromatography coupled time-of-flight mass spectrometry (UPLC-ESI-QTOF/MS) profiling, we further quantified the significantly increased intratumoral S1P in SPF mice treated with Capecitabine and Nivolumab compared with SPF mice of saline group or GF mice (Fig. EV2C). This microbial pattern was confirmed by Sankey network of integrative correlation analysis that *B. fragilis* was closely associated with up-regulated intratumoral S1P in Capecitabine and Nivolumab combination treatment group (Fig. EV2D). Pearson correlation coefficient between major classes of microbial SLs and candidate taxa (at the species level) also manifested significantly negative correlation of *B. fragilis* with intratumoral microbiota-derived S1P(d17:1) or S1P(d18:1), respectively (*P* < 0.001, Fig. 3B). Phylogenetically, the majority of bacterial species cannot produce sphingolipids. But *Bacteroides*, one of the most predominant commensal genera in intratumoral microbiota, can produce and provide a source of ceramides with both odd (C17:0) and even (C18:0) numbers of hydrocarbons.(Brown *et al*, 2019; Sepich-Poore *et al*, 2021) The biosynthesis of intratumoral S1P originated from the phosphorylation of sphingosine *via* enzymes microbial sphingosine kinase 1/2 (SphK1/2, Fig. EV3A). To identify key enzymes of modulating downstream S1P levels in response to pharmacological agents, we biosynthesized mutant bacterial strains with inactivated SphK1/2 (BFΔSphK1 and BFΔSphK2, Fig. EV3B). Then we co-cultured SW620, SW480, HT-29, HCT116 or HCT-15 cells with wild-type *B. fragilis* (BFWT) or BFΔSphK1/2 cells within the transwell system, and added ceramide to determine whether *B. fragilis-*derived S1P can be transferred to mammalian hosts to interfere with drug efficacy (Fig. EV3C). High-resolution MS2 spectra also revealed the significantly decreased S1P in various CRC cell lines when co-cultured with BFΔSphK2, in comparison to those co-cultured with BFWT or BFΔSphK1 (Fig. EV3D). These indicated that S1P catalyzed by microbial SphK2 can penetrate into CRC cells, thus improving the combination treatment of Capecitabine and Nivolumab.

**Figure 3.**
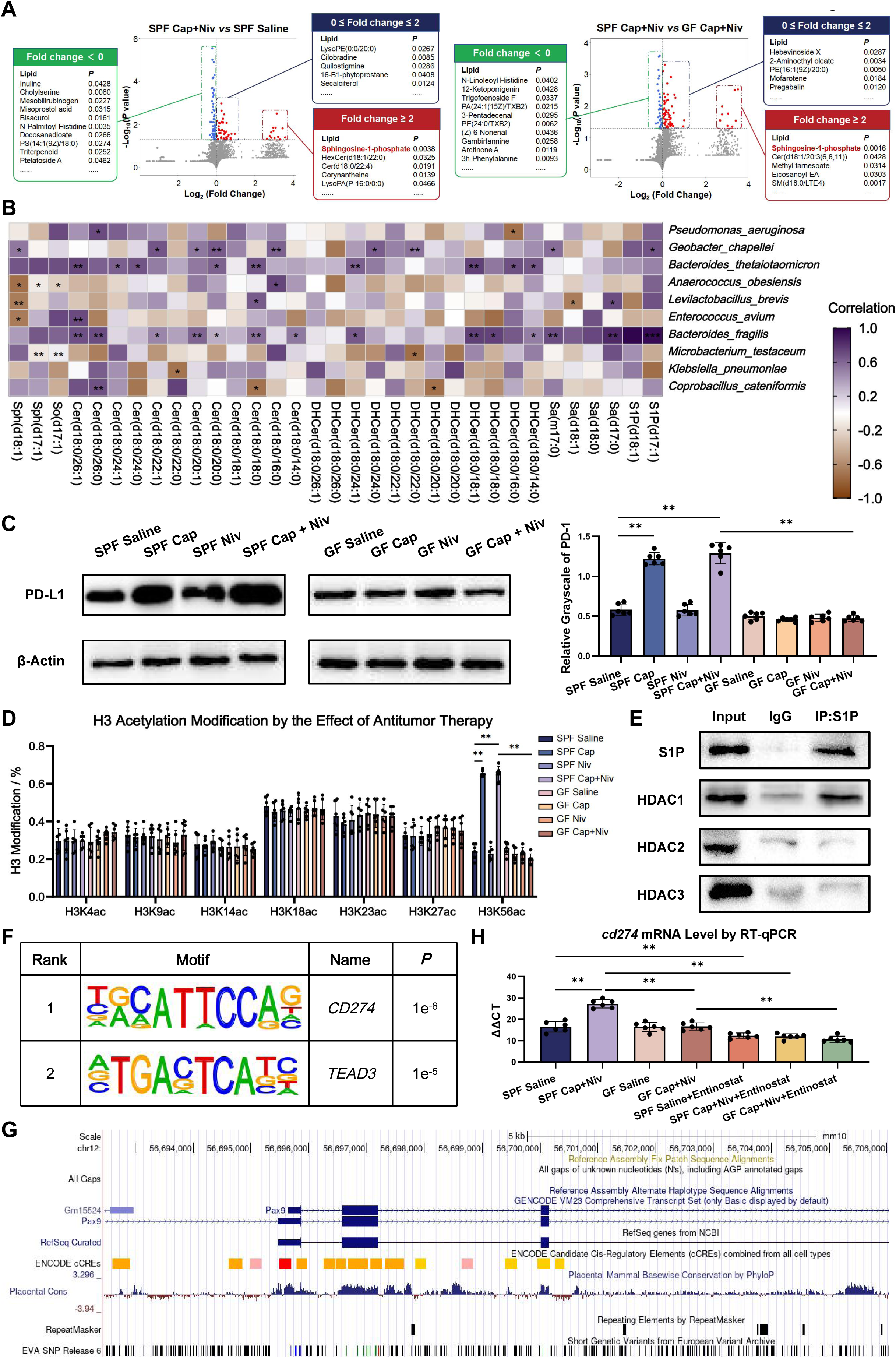
*B. fragilis*-derived sphingosine-1-phosphate (S1P) upregulated the histone deacetylation of H3K56. **(A)** Volcanic plots to illustrate intratumoral metabolites difference between Capecitabine-Nivolumab-treated SPF mice and saline-treated SPF mice (left panel)/ Capecitabine-Nivolumab-treated GF mice (right panel). Dots corresponding to significant lipids (*P* < 0.05, Student’s *t*-tests) were colored, in which lipids with increased fold change were colored as red, and sphingolipids with decreased fold change pertained to green. **(B)** Pearson correlation heat-map between intratumoral microbial species abundance and differential sphingolipids after Capecitabine-Nivolumab treatment. **(C)** The comparisons of PD-L1 levels across groups by western blotting assay. Data are means ± *SD*, \**P* < 0.05, \*\**P* < 0.01 (Student’s *t*-tests). **(D)** Genome-wide screening assay analyzed the levels of acylation sites of H3 in the nucleus of colorectal epithelial cells isolated from SPF and GF mice. **(E)** Co-IP assay showing an interaction of S1P with HDAC1 in colorectal cancer cells. **(F)** DNA motif analysis in ChIP-seq peaks of H3K56ac showing the significant enrichment of *CD274* motif (Hypergeometric test). **(G)** UCSC Genome Browser view of H3K56ac-binding sites of *CD274* for SPF mice treated with Capecitabine-Nivolumab combination. **(H)** The mRNA level of the fragment chr3:56,696,700-56,696,900 analyzed by RT-qPCR assay in each group.

It was previously reported that S1P serves as an endogenous inhibitor of HDAC1/2 to regulate histone acetylation modification in *p21* and *c-fos* promoter region.(Hait *et al*, 2009) However, the specific modification sites by the combination treatment of Capecitabine-Nivolumab and whether it affects the transcription of other target genes are still unclear. Considering that Nivolumab efficacy was mainly affected by the expression of intratumoral PD-L1, we thus determined the levels of intratumoral PD-L1 under different drug exposures, pinpointing higher intratumoral PD-L1 in SPF mice than those of GF mice (Fig. 3C). Extracting total histone from their CRC tissues, we determined the main differences in histone modification sites of SPF mice treated with Capecitabine or Capecitabine-Nivolumab combination as H3K56 acetylation (H3K56ac), comparing with other groups (Figs. 3D and EV4A). To further confirm the mechanism of S1P affecting H3K56ac, co-immunoprecipitation (Co-IP) was applied to reveal that S1P directly bound to HDAC1 rather than HDAC2/3, GCN5, HAT1, PCAF, HBO1 or p300/CBP (Figs. 3E and EV4B). We also performed glutathione S-transferase (GST) pull-down to validate the direct interaction between S1P and HDAC1 *in vitro* (Fig. EV4C), which suggested that bacteria-derived S1P binding to intratumoral HDAC1 may function as the key mediator of Capecitabine-Nivolumab combination treatment. Then RNA sequencing (RNA-seq) was applied for CRC tissues of SPF and GF mice treated with Capecitabine and Nivolumab, which resulted into a total of 1,199 differentially expressed genes (DEGs), including 593 up-regulated and 606 down-regulated genes (*P* < 0.05, Fig. EV4D). To verify whether the combination treatment of Capecitabine and Nivolumab could prompt epigenetic changes within CRC cells, chromatin immunoprecipitation sequencing (ChIP-seq) analysis for H3K56ac was thus performed in CRC tissues of SPF mice and GF mice. There were 9,353 (39.8%) gained and 5,946 (25.3%) lost H3K56ac peaks in SPF mice compared with GF mice, but their distributions of H3K56ac across the genome were similar (Fig. EV4E). As compared with GF mice, the binding sites of H3K56ac and downstream target genes in SPF mice were more concentrated in the promoter region and exon region. We subsequently sought to identify transcription factors involved in chromatin remodeling by Capecitabine-Nivolumab combination treatment in SPF mice. Motif analysis of H3K56ac peaks in SPF mice using HOMER revealed that *CD274* motif was significantly enriched at H3K56ac peaks (Fig. 3F). Using UCSC database, we ascertained the binding sequence of H3K56ac and PD-L1 coding gene to locate in the exon region of chr3:56,696,700-56,696,900. The presence of microbial S1P significantly activated the transcription of the above PD-L1 synthetic gene by inhibiting tumoral HDAC1 activity and increasing the acetylation level of H3K56 (Fig. 3G). Primers for qPCR amplification were then designed for chr3:56,696,700-56,696,900 sequence in CRC tissues treated with Capecitabine-Nivolumab combination and their control counterparts. SPF mice receiving combination therapy manifested the up-regulated transcription level of chr3:56,696,700-56,696,900. Entinostat, the inhibitor of HDAC1, effectively suppressed the transcription level of this segment (*P* < 0.01, Fig. 3H). It thus suggested that Capecitabine in combination therapy promoted the production of microbial S1P, and synergistically enhanced the efficacy of Nivolumab by activating PD-L1 expression in CRC tissues.

### Intratumoral SphK2 activity of *B. fragilis* modulated the anti-tumor efficacy of Capecitabine-Nivolumab combination therapy

To tightly monitor microbial SphK2-catalyzed S1P production, we mono-colonized 6-week-old GF mice with BFWT or BFΔSphK2 strain (Fig. EV5A), prior to the above treatments (Fig. 4A). As shown in Fig. 4B, both Capecitabine and Nivolumab obviously affected the body weights of GF mice colonized with BFWT (GF^BFWT^) and BFΔSphK2-colonized GF mice (GF^BFΔSphK2^). The combination of Capecitabine and Nivolumab induced greater weight gain in GF^BFWT^ mice than those of GF^BFΔSphK2^ mice (*P <* 0.01). Capecitabine outperformed Nivolumab in ameliorating diarrhea, bloody feces and tumor proliferation of both GF^BFWT^ and GF^BFΔSphK2^ mice, while symptoms of GF^BFWT^ mice were superior to GF^BFΔSphK2^ mice when treated with Capecitabine-Nivolumab combination therapy (*P <* 0.01, Figs. EV5B,C and 4C). GF^BFΔSphK2^ mice treated by Capecitabine/Nivolumab manifested no distinct difference in colonic length, tumor number and volume, compared with GF^BFWT^ mice (Figs. 4D,E and EV5D). Nevertheless, GF^BFWT^ mice treated with Capecitabine-Nivolumab combination therapy showed statistically longer colon length, less tumor number and volume than those of GF^BFΔSphK2^ mice (*P <* 0.05; Figs. 4D,E and EV5D). High-resolution MS2 spectra quantified the dramatic increase of intratumoral S1P in GF^BFWT^ mice treated with Capecitabine alone or combination therapy, as compared with the saline controls (*P <* 0.01, Fig. 4F). Likewise, western blot assay also showed significantly increased PD-L1 expression in Capecitabine alone or combination treatment groups (*P <* 0.01, Fig. 4G). High levels of intratumoral PD-L1 were similar between GF^BFWT^ and GF^BFΔSphK2^ mice treated with Capecitabine, but significantly differentiated between GF^BFWT^ and GF^BFΔSphK2^ mice treated with saline (Fig. 4G), confirming the microbial SphK2-mediated synergistic effects of Capecitabine and Nivolumab.

**Figure 4.**
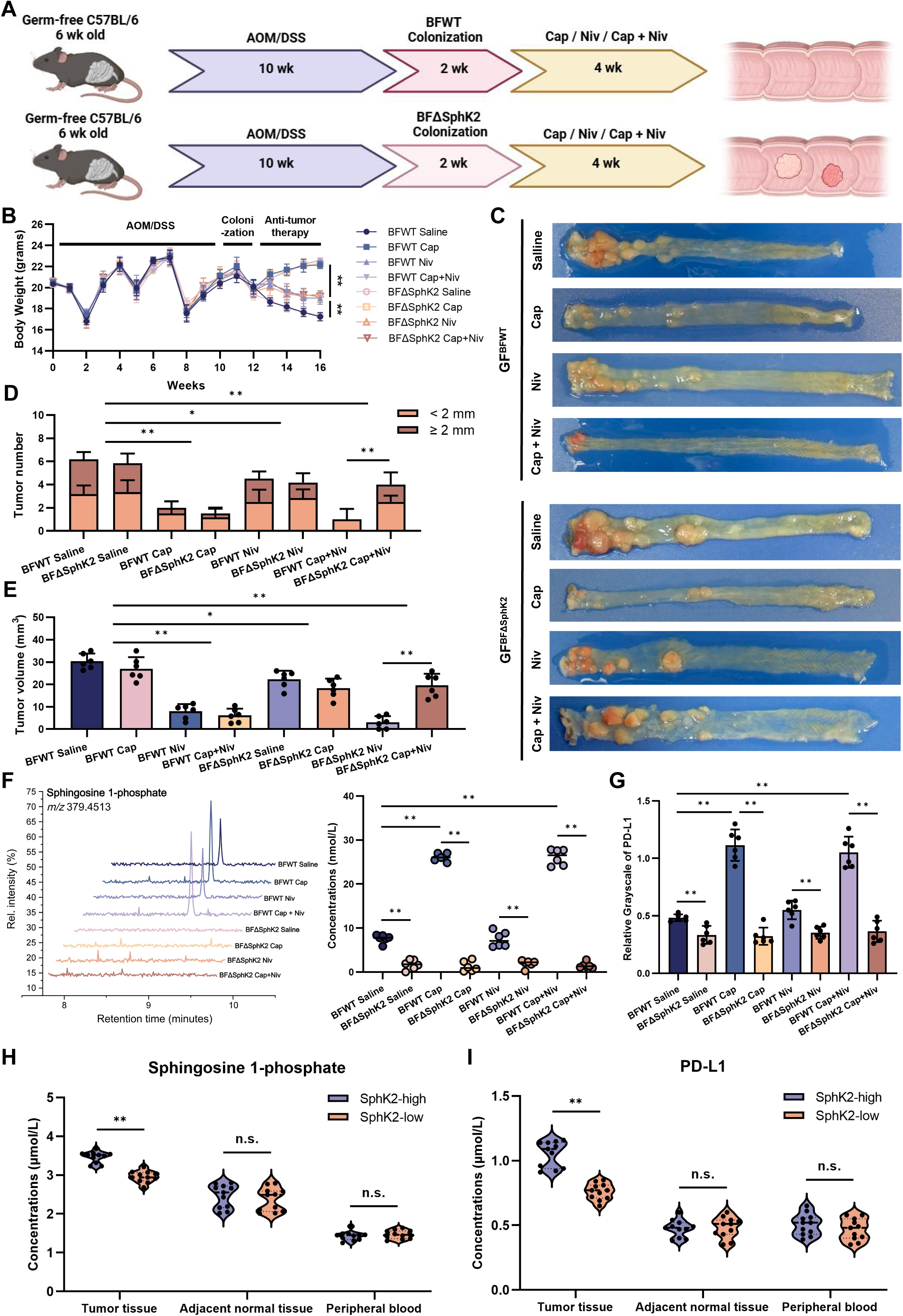
S1P-production capacity of colonized *B. fragilis* affected efficacy of Capecitabine-Nivolumab combination. **(A)** Experimental setting: age-matched GF mice were simultaneously induced with 10 weeks of AOM/DSS before 2 weeks of BFWT or BFΔSphK2 colonization, followed by the treatment of saline, Capecitabine, Nivolumab or their combination for 4 weeks. **(B)** Body weight of each group during 16 weeks of AOM/DSS induction and 2 weeks of BFWT/BFΔSphK2 colonization, then treated with anti-tumor therapies for 4 weeks. *n* = 6 mice per group, \**P* < 0.05, \*\**P* < 0.01 (Student’s *t*-tests). **(C)** Colorectal longitudinal sections of each group. **(D,E)** The number **(D)** and volume **(E)** of tumors in the colon and rectum were measured in GF^BFWT^ and GF^BFΔSphK2^ mice after the above 16 weeks of CRC model construction and treatments. **(F)** LC-MS/MS to quantify intratumoral S1P across the groups, \*\**P* < 0.01 (Student’s *t*-tests). **(G)** Differential expressions of intratumoral PD-L1 in GF^BFWT^ and GF^BFΔSphK2^ mice across groups. Data are means ± *SD*, \*\**P* < 0.01 (Student’s *t*-tests). **(H)** Quantification of intratumoral S1P concentrations, adjacent and peripheral tissues of patients with high or low microbial SphK2 activity, *n.s.* > 0.05, \*\**P* < 0.01 (Student’s *t*-tests). **(I)** Comparisons of intratumoral PD-L1 levels across CRC patients by WB assay. Data are means ± *SD*, *n.s.* > 0.05, \*\**P* < 0.01 (Student’s *t*-tests).

### Single-cell dynamics and cross-tissue landscape of CRC patients treated with Capecitabine-Nivolumab combination therapy

To elucidate the dynamic effects of Capecitabine combined with Nivolumab treatment, we applied single-cell RNA sequencing (scRNA-Seq) to longitudinally track CRC patients receiving this combination therapy throughout the treatment course. Based on the strict inclusion criteria, 22 CRC patients were ultimately enrolled with each subject undergoing baseline (pre-treatment) sampling followed by one or more post-treatment samplings after initial drug administration, including the follow-up surgery when necessary (Fig. 5A). To comprehensively map the local and systemic dynamics, CRC tissues obtained from colonoscopic or surgical biopsies were paired with peripheral blood and adjacent normal tissue samples, which were all processed immediately for matched scRNA-Seq (Fig. 5B). We divided all subjects into SphK2-high activity group and SphK2-low activity group based on their microbial SphK2 activities, and examined 918,367 cells for further analysis according to the parameter settings (nFeature-RNA > 500 and percentage. mt < 20; Fig. 5C). The transcriptomes of 815,265 high-quality single cells were selected after stringent quality control and filtration. Unsupervised clustering combined with classical marker gene annotation identified six major cell populations: T cells, B cells, innate lymphoid cells (ILCs), myeloid cells, stromal cells and epithelial cells (Fig. 5D). Separated sub-clustering on each major compartment resulted into 63 fine-grained cell subsets, which were all characterized by their distinctive expression profile as well as distribution patterns across the blood, adjacent normal tissues and tumor tissues (Fig. 5E). To statistically quantify tissue enrichment of each sub-population, we measured the ratio of observed to expected cell numbers (Robs/exp). By comparing organizational distribution preferences, we found that helper T cells (c02-CD4_CTSH), memory B cells (c20uCD27), exhausting T cells (c03_CD8_LAYN), macrophages (c29_Sph_SCGR3A) and effector memory T cells (c08_CD8_GZMK) were mainly enriched in CRC tissues (Fig. 5F). For each major and fine-grained cell type, the baseline cellular abundance and cellular dynamics (characterized as the difference between the post-treatment and baseline abundance) were calculated. Intratumorally, we observed disparate response associations of the four major immune cell types. Notably, B cells, ILCs and myeloid cells demonstrated no significant inter-group difference from baseline to post-treatment. After combination therapy, the SphK2-high activity group manifested significantly elevated T cell infiltration when compared to the SphK2-low group (Fig. 5G), suggesting enhanced T cell-mediated anti-tumor immunity in the former group. Metagenome profiles of CRC patients also indicated similar α-diversity, β-diversity and dominant colonies between two groups (Fig. EV5E-H). By collecting their CRC tissues, we determined SphK2-high patients with markedly higher intratumoral S1P and PD-L1 concentrations (*P <* 0.01, Fig. 4H,I). It thus provides us with critical insights into the potential role of microbial SphK2 in modulating immunotherapy response and offers novel perspectives for optimizing personalized strategies of Capecitabine and Nivolumab.

**Figure 5.**
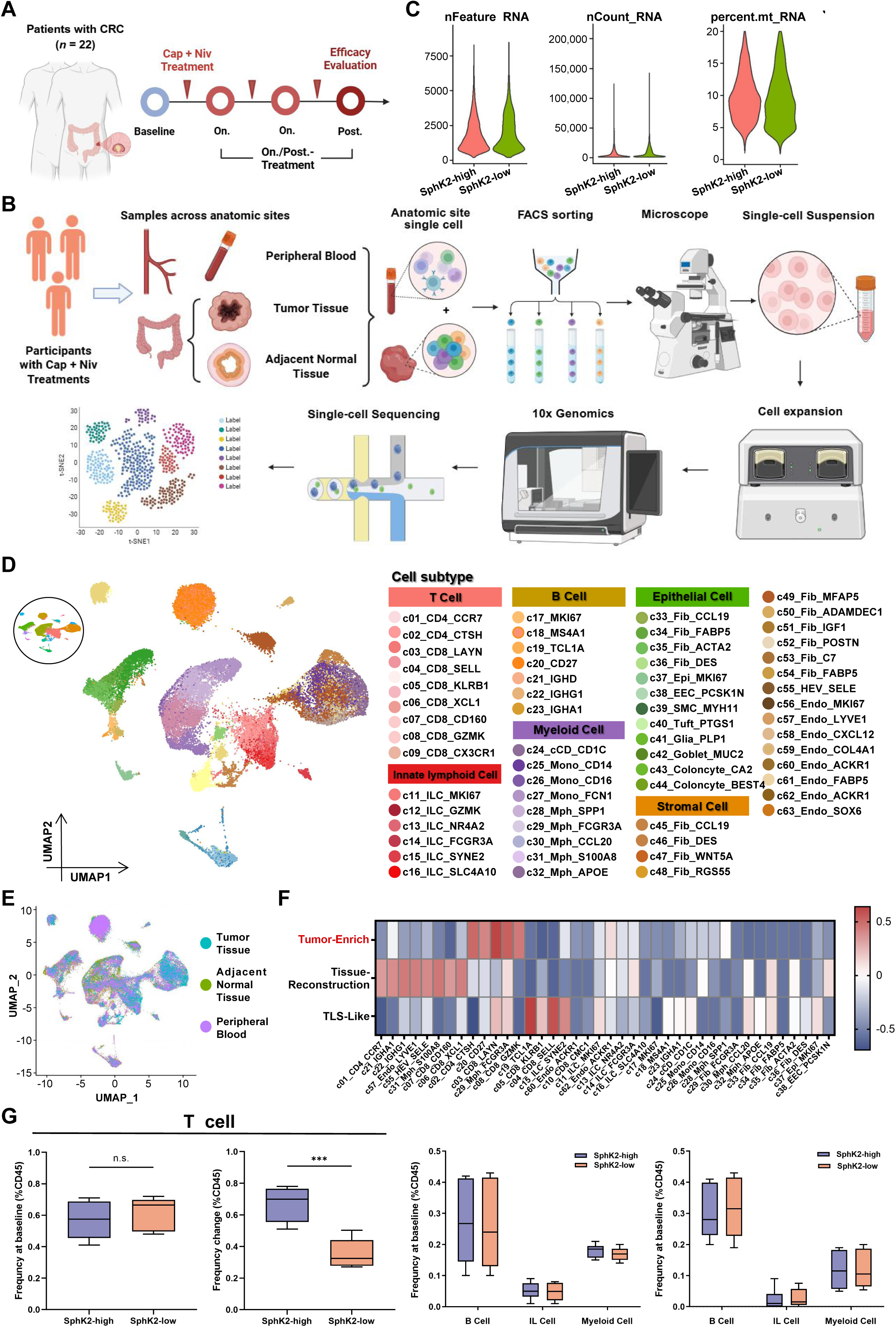
Dynamic single-cell landscape of CRC patients during Capecitabine-Nivolumab treatment. **(A)** Dynamic tracking of 22 CRC patients undergoing Capecitabine-Nivolumab treatment. **(B)** Workflow of sample collection and analysis of single-cell RNA sequencing. **(C)** Quality control of scATAC-seq data (left panel). Distribution of accessible chromatin regions per nucleus (nFeature; cutoff: > 500; middle panel). Log_10_-scaled distribution of total fragments per nucleus (nCount; right panel). Proportion of mitochondrial reads (percent.mt; exclusion threshold: < 20%). **(D)** UMAP plot of cell clusters. Cell types were identified by the regulation activity matrix and then visualized by UMAP. Cells were colored according to annotated cell types. **(E)** UMAP plots showing the anatomic site. **(F)** Tissue preference of each cluster estimated by observed-to-expected ratio (Robs/exp). The Robs/exp was *Z*-score transformed. **(G)** Box-plots showing the relative expression levels of immune cells in 22 CRC patients. *P*-value was calculated using Student’s *t*-test, \*\*\**P* < 0.01.

### Associations of exhausted T (Tex) cells with the combination efficacy of Capecitabine and Nivolumab

Considering the aforementioned findings that microbial SphK2 mediated Capecitabine-Nivolumab combination efficacy, coupled with the observed dynamic patterns of diverse immune cell populations, we subsequently focused on the T-cell compartment and sub-clustering T cells into their established phenotypes using a graph-based clustering algorithm (Fig. 6A). Unsupervised clustering analysis of all T-cell sub-clusters revealed that exhausted T cells constituted the predominantly altered sub-population (Fig. 6B). Each population was defined by known classical signature genes. Given that Tex cells exhibited significant expression of *PDCD1* and other exhaustion related genes, including *LAYN*, *HAVCR2* and *CXCL13*, we thus directed our focus to *CD8*^+^ Tex (Fig. 6C,D). Longitudinal tracking of exhausted T cell sub-population dynamics throughout the treatment course further substantiated this distinct phenotypic conversion between SphK2-high and SphK2-low cohorts, which suggested that Capecitabine combined with Nivolumab may reverse T cell exhaustion by inhibiting *LAYN* expression. To decipher the pertinent effects of Capecitabine combined with Nivolumab blockade among deferentially responsive groups, we performed comprehensive profiling of exhausted T cell sub-populations to delineate distinct patterns between SphK2-high and SphK2-low cohorts. Notably, post-treatment analysis revealed the substantial accumulation of terminally exhausted T cells (c03_CD8_LAYN) in CRC tissues from SphK2-low patients. Intriguingly, the abundance of terminally exhausted and proliferating exhausted T cell populations was predominantly replaced by effector memory T cells (c08_CD8_GZMK) following immunotherapy for microbial SphK2-low patients (Fig. 6E). Gene-set enrichment analysis (GO-BP and Hallmark) identified the significant up-regulation of immune response pathways in SphK2-high activity group following Capecitabine and Nivolumab treatment (Figs. 6F and EV6). Building upon the observed dynamic interplay between *LAYN^+^* exhausted *CD8^+^* T cells and functionally competent T cell populations, we comprehensively evaluated their combined prognostic significance in our patients’ follow-up. Notably, treatment with Capecitabine and Nivolumab demonstrated superior clinical outcomes in microbial SphK2-high patients, with significantly improved disease-free survival (DFS: 70 months *versus* 130 months; HR 0.81, *P* < 0.05) and overall survival (OS: 180 months *versus* 230 months; HR 0.69, *P* < 0.05) compared to microbial SphK2-low counterparts (Fig. 6G). Predicting which patients will benefit from preoperative Capecitabine and Nivolumab, is the crucial objective to improve patient outcomes and to provide organ-preserving options for operable CRC. Based on the above indicators treated with Capecitabine and Nivolumab, the presence of microbial SphK2 activity and the relative abundance of intratumoral *B. fragilis* were negatively correlated with TMB and CEA levels (Fig. 6H).

**Figure 6.**
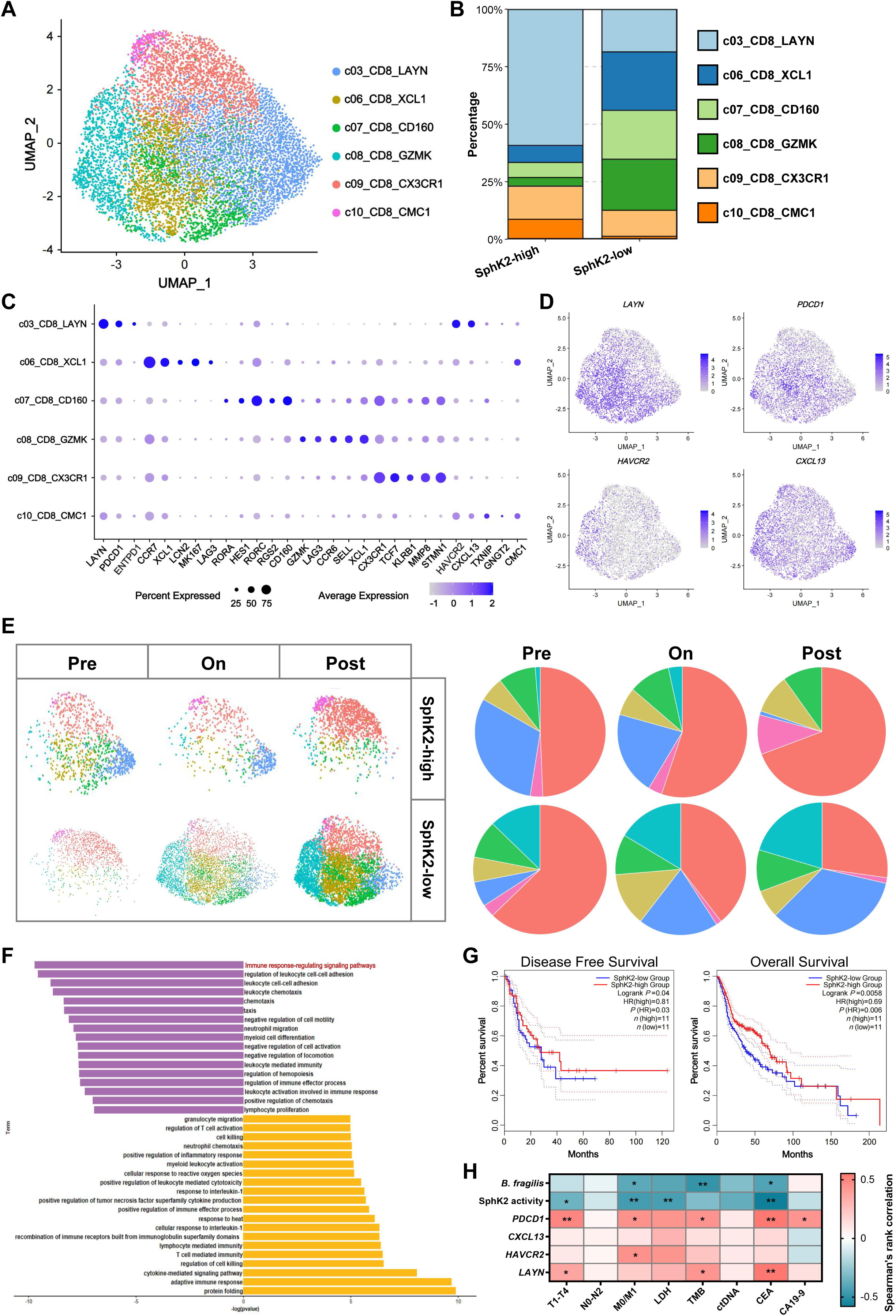
Tumor-reactive-like *CD8*^+^ T cells under Capecitabine-Nivolumab treatment. **(A)** Sub-clustering of *CD8^+^* T cell clusters, colored and labeled by sub-types. **(B)** Clustering of *CD8*^+^ T cell clusters in groups with different microbial SphK2 activities. **(C)** Bubble heat-map showing the expression of representative signature genes of *CD8*^+^ T sub-types. Both the color and size indicate scaled mean expression across cell sub-types for each gene. **(D)** UMAP plots showing expression of representative genes. **(E)** UMAP plots showing the dynamics of *CD8*^+^ T cell clusters following the Capecitabine-Nivolumab treatment (left panel). Pie charts showing the *CD8*^+^ T sub-type composition of representative changes across different treatment stages (right panel). **(F)** GO analysis analyzes pathway enrichment after the treatment of Capecitabine-Nivolumab combination. Microbial SphK2-high expression group is indicated in purple; pathways associated with inflammation with low microbial SphK2 expression are indicated in yellow. **(G)** The Kaplan-Meier overall survival curve in patients with CRC, grouped by different expression levels of microbial SphK2. Patients separated by high (*n* = 11) or low-level (*n* = 11) microbial SphK2 signature exhibit different disease-free survival and overall survival *(P* < 0.05). **(H)** Microbial SphK2 activity, relative abundance of intratumoral *B. fragilis*, expressions of *PDCD1*, *CXCL13*, *HAVCR2*, *LAYN* to correlate with CRC indicators.

## Discussion

Intratumoral microorganisms primarily originate from intestines, oral cavities and adjacent tissues, where intestinal mucosal damage and blood circulation facilitate their colonization within tumors.(Bertocchi *et al*, 2021; Xie *et al*, 2022; Yang *et al*., 2023) Predominant microorganisms enriched in CRC include *Fusobacterium nucleatum,* polyketide synthase (pks)^+^ *Escherichia coli* and *B. fragilis.*(Cao *et al*, 2021; Consortium, 2020; Hong *et al*, 2021) These intratumoral microbiota promote CRC progression by modulating intestinal epithelial cells, neoplastic cells and tumor microenvironment. Mechanistically, their oncogenic effects involve critical biological processes such as genotoxic DNA damage induction, dysregulation of apoptotic pathways and promotion of epithelial-mesenchymal transition.(Liu *et al*, 2021; Qu *et al*, 2023) The intricate interplay between intratumoral microbiota and host anti-tumor immunity provides a nuanced perspective for developing combination therapies, particularly involving agents such as Capecitabine and Nivolumab. In this study, we have conducted a comparative evaluation of the anti-tumor efficacy between GF and SPF mice models under Capecitabine plus Nivolumab combination therapy. It manifested significantly superior tumor inhibition in SPF mice as compared to their GF counterparts, indicating a pivotal regulatory role of intratumoral microbiota in therapeutic responsiveness. This observation aligns with the established literature documenting that GF or antibiotic-treated mice exhibited impaired responses to immune-checkpoint blockade therapy, whereas *Bacteroides* supplementation effectively restores therapeutic efficacy.(Vétizou *et al*, 2015) Through intratumoral microbiota composition and metabolic profiles across treatment groups, we identified intratumoral microbiota-derived S1P as the critical metabolite in mediating differential therapeutic outcomes. Of note, elevated intratumoral S1P derived from *B. fragilis* directly interacts with histone deacetylase HDAC1 in CRC cells, suppressing its deacetylase activity and enhancing H3K56ac (Hait *et al*., 2009). Through ChIP-seq and ChIP-qPCR analyses, we further demonstrated that intratumoral S1P-mediated H3K56ac exhibits significant enrichment within the *CD274* gene locus, leading to the markedly up-regulated PD-L1 transcription in CRC tissues of SPF mice subjected to combination therapy. These findings collectively indicate that microbial S1P enhanced CRC tissue dependency on the PD-L1 pathway by inhibiting HDAC1 and elevating histone acetylation at the PD-L1 genomic region, ultimately synergizing with Nivolumab to potentiate therapeutic efficacy.

Previous studies have revealed that deficiency of *Bacteroides*-derived sphingolipids induces intestinal inflammation and disrupted host lipid metabolism in GF mice,(Brown *et al*., 2019) underscoring the pivotal role of microbiota-synthesizing S1P to regulate host immunoregulation processes. Our single-cell sequencing analyses further elucidated the relationship between microbial SphK2 activity and dynamics of tumor-infiltrating T cell lineages. In patients with high microbial SphK2 activity, combination therapy induced a significant expansion of effector memory *CD8^+^* T cells within the tumor microenvironment, accompanied by a relative reduction in exhausted T cell subsets. In contrast, patients with low microbial SphK2 activity retained a higher proportion of terminally exhausted T cells. Gene set enrichment analysis demonstrated pronounced enrichment of immune activation signaling in the SphK2-high activity cohort post-treatment. Furthermore, our data suggest that microbial SphK2 may regulate the equilibrium between T cell exhaustion and memory differentiation through analogous mechanisms, thereby dynamically modulating immune response intensity. Integrated analysis of clinical cohort data revealed that patients with elevated microbial SphK2 activity exhibited superior clinical outcomes following Capecitabine plus Nivolumab therapy, including significantly prolonged disease-free survival (DFS) and overall survival (OS). These findings not only provide a novel biomarker for clinical stratification but also indicate microbial sphingolipid metabolites as potential predictive tools for assessing patient responsiveness to immunotherapeutic interventions.

Taken together, our systematic investigation elucidates that intratumoral S1P produced by *B. fragilis* modulates immune checkpoint expression through an epigenetic regulatory pathway, potentiating the therapeutic efficacy of Capecitabine and Nivolumab combination therapy. This discovery significantly enhances our comprehension of intratumoral microbiota involvement in oncotherapy and immune regulation. Future research should prioritize delineating mechanistic roles of other pivotal microorganisms and their metabolites in cancer treatment paradigms, and translating these findings into clinically actionable intervention strategies would enable more personalized and precision therapeutic approaches for CRC and other malignancies.

## Methods

### Animal experiments

All procedures involving animals were carried out according to the protocols approved by Peking University First Hospital Experimental Animal Center (Ethical No. 2022-23142). C57BL/6J germ-free (GF) or specific pathogen-free (SPF) mice were housed at controlled temperature (20-22°C) and humidity (40-60%) with a 12-h light-dark cycle and access to food and water ad libitum, which were acclimatized for one week before grouping. CRC mice model was established by intraperitoneal injection of 10 mg/kg azoxymethane (AOM) once, followed by three cycles of dextran sulfate sodium (DSS) administration (2% DSS in drinking water for 1 week and normal drinking water for 2 weeks per cycle). For the mice experiments in Figs. 1 and 2, GF and SPF mice were treated with varieties of CRC therapy for 4 weeks (*n* = 6 each group). At the end of week 14, all mice were euthanized by cervical dislocation. The colorectal tract was excised from the cecum to the anus, longitudinally incised along its length, photographed, and subjected to tumor enumeration. The number of colorectal tumors was recorded and CRC tissues were collected for further analysis (refer to ‘Metagenomic sequencing and analysis’ and ‘Metabolome analysis of intratumoral microbiota’ sections). Subsequently, quantitative PCR (qPCR) was applied to validate microbial abundances, and LC-MS was used to quantify metabolites of the initial screening results. After false discovery rate (FDR) correction, statistical significance was assessed using Wilcoxon’s rank sum test with a *P*-value cut-off of 0.05. For *in vivo* experiments depicted in Fig. 4, all mice also initially received AOM combined with DSS to induce CRC, and divided into four treatment groups: 1) Saline group 2) Capecitabine group 3) Nivolumab group and 4) Capecitabine-Nivolumab combination group. Prior to 2-week drug administration, either BFWT or BFΔSphK2 strains were colonized through dietary supplementation for 4 weeks.

### Metagenomic sequencing and analysis

Sequences were grouped into operational taxonomic units (OTUs) using UPARSE (v.7.0.1001), clustering all high-quality tags from each sample at a 97% similarity threshold. Taxonomic annotation and statistical evaluations at various classification levels were carried out against the SILVA138 database. Alpha diversity indices (ACE, Chao1, Shannon and Simpson) were used to measure community richness and diversity of microbe. For beta diversity analysis, uniFrac distances were computed *via* QIIME (v.1.7.0), and UPGMA (Unweighted Pair Group Method with Arithmetic Mean) clustering trees were generated to visualize sample relationships. Principal component analysis (PCA) and principal coordinates analysis (PCoA) were performed in R software to reveal sample-to-sample variation, using the FactoMineR and ggplot2 packages to reduce dimensions of raw normalized variables. Potential biomarkers were identified through Linear Discriminant Analysis Effect Size (LEfSe), focusing on statistically significant (*P* < 0.05) and biologically relevant (LDA score ≥ 4) features to identify taxa with notable inter-group differences.

### Metabolome analysis of intratumoral microbiota

A total of 50 mg of CRC tissue was collected from each mouse for metabolomic profiling. For untargeted analyses, samples were initially treated with 80% cold methanol. Following centrifugation at 21,500 *g* for 15 minutes at 4 °C, the supernatant was obtained for LC-MS evaluation. LC-MS was performed on an Agilent 1290 Infinity UPLC system (Agilent Technologies, Santa Clara, CA) in conjunction with Sciex TripleTOF 6600 mass spectrometer (Q-TOF, AB Sciex, Toronto, Canada). Chromatographic separation was carried out using a Waters BEH Amide column (2.1 mm × 100 mm, 1.7 μm). All raw MS data were transformed to mzXML format *via* ProteoWizard, then processed in R using XCMS package (v.3.2) to produce a data matrix encompassing retention time (RT), mass-to-charge ratio (*m/z*) and peak intensity. The CAMERA package (v.16.4) in R was utilized to annotate peaks, and metabolites were identified by referring to the MS2 database. The resulting metabolite list and their abundances were examined with the MetaboAnalyst package (v.4.0). Metabolites displaying significant alterations were identified using the Wilcox test, with adjusted *P* < 0.05 indicating statistical significance. Partial spearman correlation was then applied to evaluate associations between differentially altered bacteria and metabolites, and heat-maps were generated using the ComplexHeatmap R package.

### RNA sequencing analysis

Total RNA was isolated and purified by TRIzol reagent following the manufacturer manual. The RNA concentration and integrity were evaluated by NanoDrop ND-1000 (NanoDrop, Wilmington, DE, USA) and Bioanalyzer 2100 (Agilent, CA, USA). Then poly (A) RNA was purified from 1μg total RNA by Dynabeads Oligo (dT) 25-61005 (Thermo Fisher, CA, USA) using two rounds of purification, which was cut into pieces using Magnesium RNA Fragmentation Module under 94 °C for 5-7 minutes. The fragmented RNA pieces were reversely transcribed into cDNA by SuperScript™ II Reverse Transcriptase and sequenced with illumina Novaseq™ 6000. Next, StringTie and edgeR (v.3.21) were used to evaluate the expression levels of all transcripts. The differentially expressed mRNAs and genes were picked with log_2_ (fold change) >1 or log_2_ (fold change) <-1 and statistical significance (*P* value < 0.05).

### Chromatin immunoprecipitation (ChIP) and ChIP-seq

ChIP assay was carried out following the instructions from the SimpleChIP^®^ Enzymatic Chromatin IP Kit. In brief, TPCs overexpressing TCF21 (TPC_NM_ ^TCF21^) were washed twice with cold PBS, treated with 1% formaldehyde for 10 minutes at room temperature to cross-link proteins and DNA, and then quenched using 125 mM glycine. Following cell lysis on ice, chromatin was extracted and subsequently sonicated into fragments averaging 150-900 bp in length. An aliquot of 20 μL of chromatin was reserved as input, and another 100 μL was used for immunoprecipitation with an anti-TCF21 antibody. An anti-IgG antibody served as the negative control. A total of 10 μg of anti-TCF21 antibody was added to each immunoprecipitation reaction before incubated overnight at 4 °C. The samples were then incubated with 30 μL of protein A beads for 2 hours. After reversing the cross-links and purifying the DNA, the immunoprecipitated DNA was quantified by real-time PCR. To prepare libraries, paired-end 150 bp reads were sequenced on the Illumina Xten platform, following the protocol of NEXTflex^®^ ChIP-Seq kit. For data analysis, low-quality reads were removed using Trimmomatic, and peak calling was conducted with MACS2 using default settings (bandwidth 300 bp; model fold 5, 50; *q* value 0.05). Peaks were assigned to the gene whose transcription start site (TSS) was closest to the peak summit.

### Microbial strains and gene-editing strategies

Isolated from the CRC tissues of enrolled participants, strains of *B. fragilis* were identified by comparing the 16S rRNA gene sequence with those in the NCBI reference database. The DNA fragments encrypting full-length microbial *SphK2* gene were cloned into pET28a vector with 6 × His tag at N-terminal end adopting standard molecular cloning procedures. Microbial *SphK2* was overexpressed in *E. coli Rosetta* (DE3), which were cultured to an OD600 of 0.6 at 37 °C. An internal fragment (290 bp) of the *SphK2* gene from *B. fragilis* was cloned into the pGERM suicide vector containing selective markers of *E. coli (bla*). The constructed vector was subsequently transformed into the conjugative *E. coli* S17 strain. Then *E. coli* S17 as donor bacteria and *B. fragilis* as receptor bacteria were co-incubated under aerobic conditions, which were transferred to BHI medium agar plates for mutant selection by gentamicin (200 μg/mL) and erythromycin (25 μg/mL). Primers targeting junction regions between pGERM and *SphK2* genes were utilized to pick resistant colonies for qPCR identification. Both wild-type and *SphK2*-depleting *B. fragilis* were grown at 37 °C in LB medium for 24 h. The bacterial culture medium was centrifuged for 10 minutes at 8,000 *g* and 4 °C, and the pellets were resuspended with oxygen-free PBS to obtain bacteria for oral administration. Colonization fitness of the two strains in mice was analyzed by 2% agarose gel electrophoresis and 3500 UV images (Fig. EV4A).

### Sample collecting and processing of scRNA-Seq

CRC tissue biopsies were collected from the enrolled participants, along with 1 mL of peripheral blood during each colonoscopy procedure. The Ethics Committees in Peking University First Hospital approved the study protocols (Ethical number: 2022-166). Written informed consents were obtained from all participants in this study. For CRC tissue samples, excised biopsies were immediately preserved in tissue preservation solution and transported on ice. After two washes, tissues were minced into 1-2 mm³ fragments and enzymatically dissociated using the gentleMACS Tumor Dissociation Kit in RPMI-1640 medium (Gibco) supplemented with 10% fetal bovine serum (FBS) for 60 minutes at 37 °C on a rotor. Dissociated cells were filtered through 100 μm SmartStrainers (pluriSelect) into RPMI-1640 medium containing 10% FBS, centrifuged at 400 *g* for 5 minutes, and pelleted. The supernatant was discarded, and cells were resuspended in 1 mL of erythrocyte lysis buffer and incubated on ice for 1 minute, followed by quenching with RPMI-1640 medium. After centrifugation for 5 minutes at 400 *g*, the cell pellet was resuspended in sorting buffer (PBS supplemented with 1% FBS). Peripheral blood samples were collected in EDTA anticoagulant tubes and cryopreserved until processing. After thawing to room temperature and thorough mixing, peripheral blood mononuclear cells (PBMCs) were isolated using HISTOPAQUE-1077 by centrifugation at 400 *g* for 30 minutes. The immune cell layer was carefully transferred to a new tube, washed twice with PBS, and subjected to erythrocyte lysis as described above. The isolated immune cells were finally resuspended in sorting buffer.

### Single-cell sources and data generation

To obtain high-quality cell pools prior to subsequent library construction, single-cell suspensions isolated from CRC tissue samples were stained with anti-7AAD antibody and anti-CD235a antibody for fluorescence-activated cell sorting (FACS) using a BD Aria III cell sorter. Live cells were enriched by gating on *7AAD⁻CD235a⁻* populations to exclude residual erythrocytes. Gated cells were sorted into 1.5 mL LoBind tubes, manually counted under a microscope, and adjusted to a concentration of 500-1,200 cells/μL. Approximately 10,000-18,000 cells were loaded into the 10 × Chromium Single Cell 5’ Library and VDJ Library Construction system, followed by standardized manufacturer protocols. Purified libraries were sequenced on an Illumina NovaSeq platform with 150-bp paired-end reads.

For blood samples, red blood cells were lysed using ACK lysing buffer according to the manufacturer’s instructions. After washing with PBS containing 2% FBS, cells were stained with surface antigen-specific antibodies at 4 °C for 30-60 minutes. For intracellular staining, cells were fixed with 4% paraformaldehyde (PFA) at 4 °C for 1 hour and permeabilized with methanol at −20 °C overnight. Cells were then blocked with Purified Rat Anti-Mouse CD16/CD32 at room temperature for 10 minutes, followed by incubation with intracellular antigen-specific antibodies at 4 °C for 1 hour. Data were acquired using a BD LSRFortessa flow cytometer and analyzed using FlowJo software.

### scRNA-seq data processing

The sequencing data were mapped to the human reference sequence (GRCh38). The raw gene expression matrix from each sample was aggregated and converted into a Seurat object *via* Seurat (v.5) package in R software. Non-immune cells with > 6000 or < 200 genes or > 40% mitochondrial genes were discarded. Immune cells with > 4000 or < 200 genes or > 25% mitochondrial genes were filtered out. To further eliminate the data of doublets, we performed the scrublet pipeline for each batch of our scRNA-seq data, which were expected to exclude doublets, and the expected_doublet_rate was set at 0.05. The gene expression matrices were normalized to the total unique molecular identifiers (UMI) counts per cell and transformed to the natural log scale. To correct the technical and biological variations and increase the accuracy of cell type designation, we applied canonical correlation analysis implemented in Seurat to all samples before cell type identification.

Seurat workflow was employed to perform dimensionality reduction and unsupervised clustering. Initially, the top 2000 highly variable genes (HVGs) were selected using the *FindVariableFeatures* function with parameters set to *method = “vst”*. Subsequently, the *ScaleData* function was applied to regress out the effects of total UMI counts and mitochondrial gene percentage from the HVG expression matrix. PCA was conducted on the scaled HVG expression matrix *via* the *RunPCA* function of Seurat, retaining the top 30 principal components for downstream analyses. A systematic two-round unsupervised clustering strategy based on the *scanpy.tl.leiden* function was executed to resolve cellular population architecture. During the first round, clusters were annotated through canonical lineage marker expression patterns, enabling identification of major cell types including T cells, B cells, ILCs, myeloid cells, stromal cells and epithelial cells. Clusters exhibiting high co-expression of two or more lineage markers were classified as doublets and excluded for subsequent analyses. Iterative refinement of major cell type annotation and doublet removal was performed to ensure purity across all primary cellular compartments. For each major cell type, a second round of unsupervised clustering was implemented using the aforementioned protocol to delineate fine-grained sub-types, with analyses exclusively restricted to the expression matrix of cells within the targeted cellular compartment. For each cellular sub-type, we assessed its distribution patterns across blood, adjacent normal tissue and tumor tissue by calculating the observed-to-expected (*Ro/e*) ratio. Pearson residuals, derived from the chi-square test, were further employed to quantify distributional differences between tumor and adjacent normal tissues. For this analysis, cells exclusively from these two tissue types were included as input for the test function.

### Statistical Analysis

At least three independent experiments were carried out, and their outcomes are shown as mean ± standard deviation (*SD*). GraphPad Prism was employed for statistical analyses. Error bars in both scatter plots and bar graphs indicate *SD*. Comparisons between groups were conducted utilizing Student’s *t*-test, Mann-Whitney U-test, one-way analysis of variance (ANOVA), Bonferroni and Scheffe post-test for continuous variables. Spearman’s Rank Correlation was used to evaluate associations between variables.

## Data availability statement

The 16S rRNA sequencing and metabolomics data supporting the results in this study are deposited in BIG Sub (Study ID: PRJCA039557) and Metabolomics Workbench (Study ID: ST003877), respectively. The minimum data-set for main figures and Supplementary Figures that support the findings of this study are openly available in Figshare (https://doi.org/10.6084/m9.figshare.29066708). Source data are provided with this paper.

## Ethics statement

All procedures involving animals were carried out according to the protocols approved by Peking University First Hospital Experimental Animal Center (Ethical No. 2022-23142). CRC tissue biopsies were collected from the enrolled participants, along with 1 mL of peripheral blood during each colonoscopy procedure. The Ethics Committees in Peking University First Hospital approved the study protocols (Ethical number: 2022-166). Written informed consents were obtained from all participants in this study.

